# neissflow: Streamlining Genomic Epidemiology of *Neisseria gonorrhoeae* with Nextflow

**DOI:** 10.64898/2026.07.22.740120

**Authors:** Kathryn Morin, Ethan Hetrick, Apurva Shrivastava, Eric Tran, John C. Cartee, Katherine Hebrank, Kim M. Gernert, Matthew W. Schmerer, Sandeep J. Joseph

## Abstract

Antimicrobial-resistant *Neisseria gonorrhoeae* (*Ng*) poses a growing global public health threat. Existing tools available for *Ng* genome analysis carry notable limitations, including incomplete resistance marker coverage, absence of species identification, and lack of phylogenetic capability. We developed neissflow, a highly parallelized Nextflow pipeline for *Ng* genome analysis that integrates five subworkflows: read preprocessing, species identification, *de novo* assembly, antimicrobial resistance (AMR) profiling, and recombination-aware phylogenetic analysis with outbreak detection. neissflow performs extensive quality control on reads, assemblies, variant calls, and phylogenetic results. Validation was performed on two datasets: a mixed-species dataset (n=158; 105 *Ng* and 53 non-gonococcal species) for sensitivity/specificity assessment, and a reproducibility dataset (n=283 replicate sequences from 17 reference strains) for consistency and phylogenetic validation. neissflow achieved 100% sensitivity and specificity for *Ng* species identification compared with MALDI-TOF and PubMLST methods. All nine AMR and typing analytes demonstrated ≥98.1% concordance with PubMLST genotype calls. Genotype-phenotype validation confirmed perfect concordance for key resistance determinants including *gyrA* mutations with ciprofloxacin resistance, 23S rRNA mutations with high-level azithromycin resistance, and *tetM* plasmid gene with high-level tetracycline resistance. Reproducibility analysis demonstrated 99.97% concordance across 3,093 analyte calls. Phylogenetic validation demonstrated 100% accuracy for both strain-level and intra-MLST clustering. neissflow is a robust, accessible, and standardized pipeline, positioning it as a valuable tool for public health laboratories engaged in *Ng* AMR monitoring and outbreak investigations.

**Importance:** *Neisseria gonorrhoeae* (*Ng*) is the second most common reported bacterial sexually transmitted infection and has developed resistance to all clinically relevant antibiotics. Surveillance of *Ng* resistance informs clinical recommendations for treatment of gonococcal infections. Whole genome sequencing (WGS) offers powerful insights into resistance mechanisms and transmission dynamics. However, many public health laboratories lack the resources needed to analyze these data effectively. We developed neissflow as an end-to-end, accessible *Ng* WGS analysis pipeline. neissflow demonstrated exceptional accuracy and reproducibility across diverse reference datasets. neissflow enables broader adoption of whole-genome sequencing-based surveillance and supports timely public health responses to emerging antimicrobial resistance in *Ng* by lowering technical barriers.

## Introduction

The rising prevalence of antibiotic resistance (AR) in *Neisseria gonorrhoeae* (*Ng*), the etiological agent of gonorrhea, poses a critical public health challenge globally (1, 2). Effective laboratory-based surveillance and timely public health response to AR *Ng* depends heavily on the rapid genetic characterization of strain types and detection of antimicrobial resistant (AMR) genetic markers (3). The expansion of whole genome sequencing (WGS) capacity in public health laboratories (PHLs) has heightened the necessity for automated tools capable of standardized and efficient genomic analysis (4–6). Automation in genomic data processing can significantly enhance the speed and reproducibility of AMR detection and strain clustering, reducing the delays often experienced with manual analyses (7). This requirement is particularly urgent due to the increasing global dissemination (8) of highly resistant gonococcal strains harboring concerning alleles such as *penA-60* (9–13) and related mosaic alleles like *penA-237* (14) associated with reduced susceptibility to ceftriaxone. These strains, frequently originating from Southeast Asian countries, are being imported into the U.S. and other regions (9–11, 14–16), amplifying the need for rapid identification and response (17). The accurate genomic analysis of these strains necessitates specialized bioinformatics support that many PHLs may lack, especially in resource-limited settings (18). The resulting gaps can hinder effective AMR surveillance and timely intervention (19).

Bioinformatics applications such as PubMLST, Gen2Epi, ARIBA, Pathogenwatch and Bactopia have been used for *Ng* genomic analysis, but each has critical limitations. PubMLST provides a web-based platform for *Neisseria* species identification, multi-locus sequence typing (MLST), and AMR marker detection (20, 21). However, PubMLST requires users to upload contigs, necessitating upstream genome assembly and quality control, and does not provide integrated phylogenetic analysis or outbreak detection. Gen2Epi, while tailored to *Ng*, focuses only on plasmid-associated AMR and sequence typing (MLST, NG-MAST, NG-STAR). Gen2Epi lacks phylogenetic analysis, automation, and requires use within a virtual machine. (22, 23). ARIBA detects known AMR markers and requires customized *Ng* reference sequences but lacks clustering or phylogenetic capabilities (24). While Pathogenwatch provides a scalable web-based platform, practical constraints remain regarding internet bandwidth requirements for raw sequence data uploads and the extent to which the platform can be tailored may pose practical limitations in local customization (25). Bactopia is a general-purpose bacterial genomic Nextflow pipeline that demands extensive user customization for *Ng* analysis (26). These limitations highlight the need for a fully automated, end-to-end pipeline for *Ng* genome analysis that integrates chromosomal and plasmid-associated AMR detection, clustering, and recombination-aware phylogenetic analysis, an essential feature given the organism’s high propensity for homologous recombination (27–31).

Bioinformatics pipelines were historically implemented as a series of scripts and system calls, resulting in complex dependencies and limited control over computational resources, which hindered portability, reproducibility, and scalability. Workflow management systems such as Nextflow (32) and Snakemake (33) address these challenges by providing high-level frameworks for modular dependency management and automated parallel execution. Nextflow is compatible with most compute environments, including High-Performance Computing (HPC) platforms and cloud services. Nextflow decouples runtime execution from pipeline logic, enabling a broad user base to run pipelines by supplying a custom configuration file. Nextflow’s modularity allows individual pipeline components to be updated or replaced without disrupting the rest of the workflow, making pipelines easier to maintain and update. Built-in dependency management leverages software containers to enforce consistent versions and configurations, preventing version conflicts and removing the burden of installation (34).

With Nextflow, we developed neissflow 2.1.0 (https://github.com/CDCgov/neissflow/), an end-to-end bioinformatics pipeline for *Ng* genomic analysis. neissflow performs comprehensive analyses; including species identification, sequence typing, AMR marker detection, and phylogenetic analysis. Additionally, it corrects for recombination during phylogenetic analysis, improving the resolution of relationships among *Ng* strains during outbreak investigations. Here we describe the development of neissflow and present validation results demonstrating its analytical performance across multiple independent datasets.

## Materials and Methods

### Development of neissflow

We developed neissflow, a Nextflow pipeline for *Ng*-specific WGS analysis. It integrates open-source tools, custom Python and Bash scripts, and Singularity containers. The pipeline logic is programmed in Nextflow Domain-Specific Language 2 (DSL2). neissflow was developed using the nf-core (v3.3.2) template, adhering to nf-core standards. The nf-core framework (34) provides standardized pipeline templates and reusable modules for common bioinformatics tools, enabling standardized, modular, and easily maintainable workflows.

neissflow accepts a sample sheet in CSV format as input containing the sample IDs and paths to FASTQ read files or FASTA assembly files. neissflow analyzes Illumina paired-end reads and/or de novo assemblies. Upon submission, the pipeline processes the input through five subworkflows. These subworkflows include 1) preprocessing raw reads, 2) species identification, 3) *de novo* genome assembly, 4) AMR typing, and 5) phylogenetic analysis. These subworkflows can also be skipped as needed by option. Quality control (QC) is performed in two stages, initially at the read level to assess sequencing quality, and subsequently at the species and assembly level to confirm *Ng* species identification and genome assembly quality. The pipeline architecture is outlined Figure 1.

**Figure 1:**
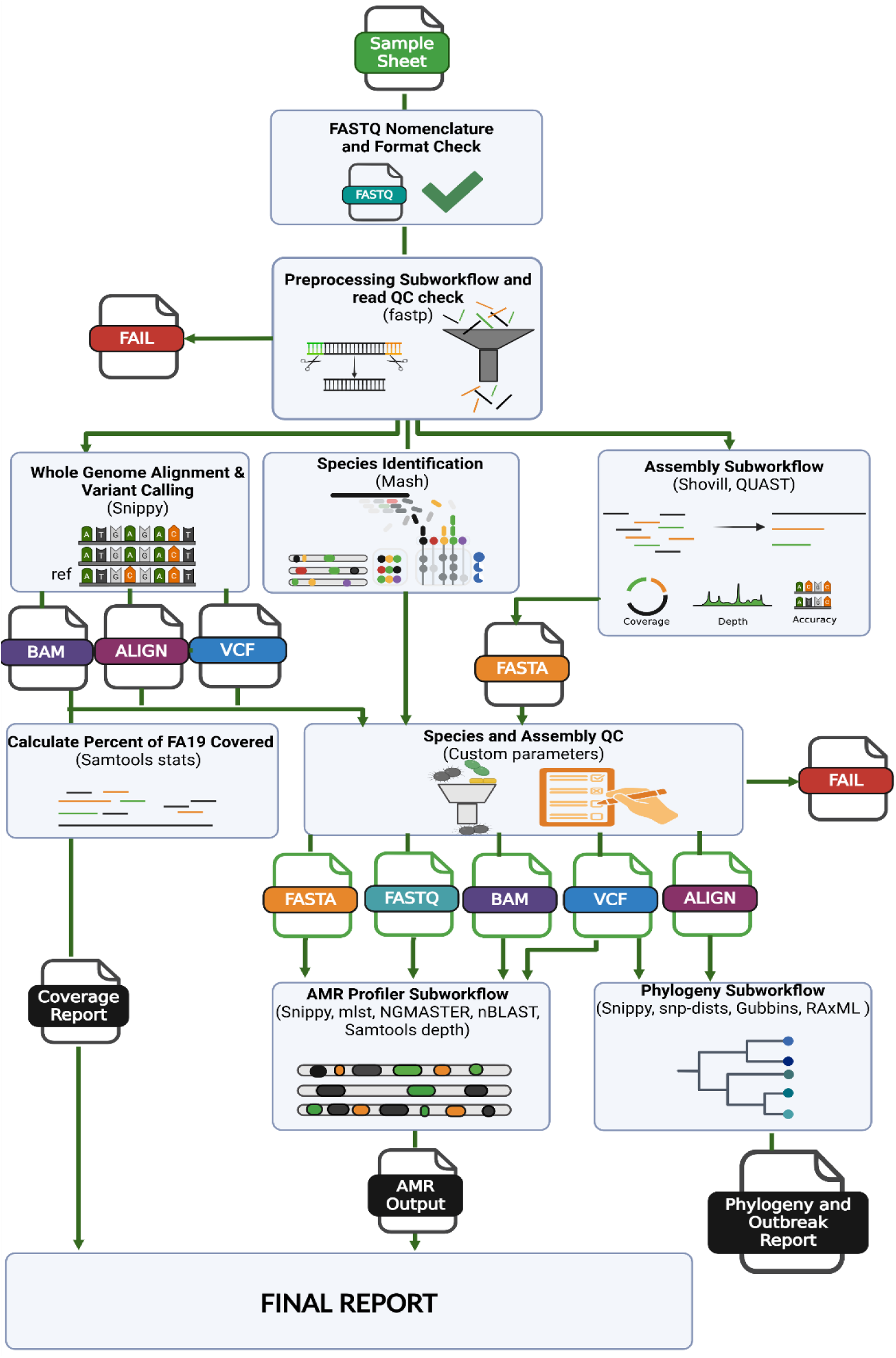
neissflow workflow diagram showing subworkflows, QC checks, and reports. Created with Bi-oRender.com. **QC** — Quality Control; **BAM** — Binary Alignment Map; **VCF** — Variant Call Format.

#### 1. Preprocessing Subworkflow and Stage 1 QC Check

The preprocessing subworkflow uses fastp (v0.23.4) (35) to perform adapter trimming and quality filtering. fastp also generates a report containing read metrics before and after processing. To pass stage 1 QC, samples must meet specific estimated depth of coverage thresholds (Table 1a) based on Illumina 2x250 paired-end sequencing.

**Table 1:**
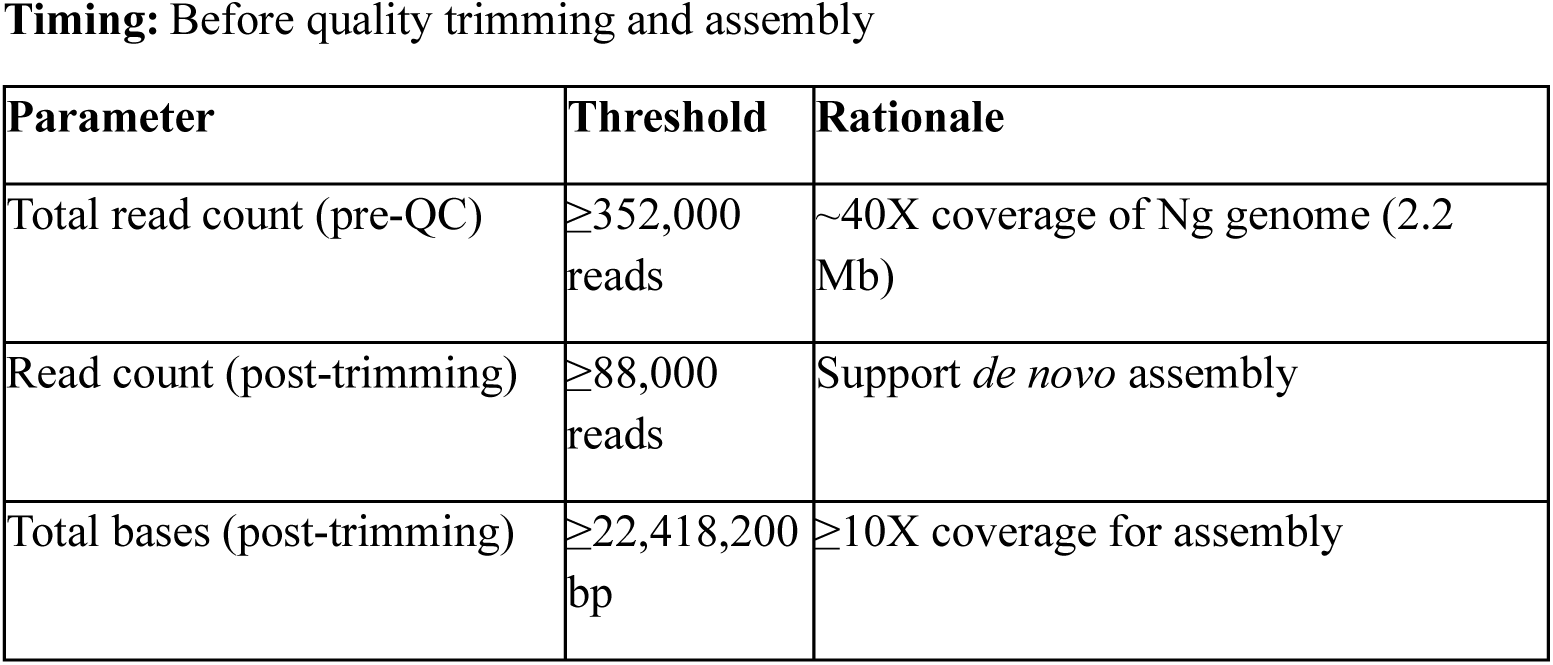
Stage 1 Quality Control - Read-level QC.

**Table 1b:** Stage 2 Quality Control - Assembly-level QC.

| Order | QC Check | Failure Condition | Note |
| --- | --- | --- | --- |
| 1 | Species Identity | Mash top hit ≠ Ng AND FA19 $>10X$ coverage $<85\%$ | — |
| 2 | Genome Coverage | FA19 $>10X$ coverage $<85\%$ | Regardless of species ID |
| 3 | Total Assembly Size | Total contig bases $>2,500,000$ bp | — |
| 4 | Assembly Contiguity & Size | Bases in contigs $>10kb$ $<1.85$ Mbp AND total bases $<2.1$ Mbp | — |
| 5 | Contiguity Fraction | Fraction of contigs $>10kb$ $<0.10$ | — |
| 6 | Mean Depth | Mean coverage $<11X$ | — |
| 7 | Joint Depth/Contiguity | Mean coverage $<15X$ AND fraction $>10kb$ contigs $<0.25$ | — |

#### 2. Species Identification Subworkflow

First, Mash (v2.3) (36) is used to screen reads against a Mash sketch of NCBI’s RefSeq database (RefSeqSketchesDefaults.msh), provided in the Mash documentation to determine the top hit (Supplementary methods). Reads are then mapped to the *Ng* FA19 reference genome (GenBank CP012026), and Samtools (v1.17) (37) stats is used to calculate the proportion of the genome covered at >10X coverage depth. Internal validation determined that high-quality *Ng* genomes consistently produced >85% coverage (Supplementary Tables S3 and S4), whereas low-quality or *Neisseria* commensal species (non-*Ng*) samples yielded <85% coverage for this metric (Supplementary table S10). This combined approach aims to exclude low-quality or non-*Ng* samples prior to downstream genomic analysis.

#### 3. Assembly Subworkflow and Stage 2 QC check

The assembly subworkflow generates *de novo* assemblies from preprocessed FASTQ files using Shovill (v1.1.0) (https://github.com/tseemann/shovill). The assemblies are then evaluated for quality. Assembly QC comprises 1) an aggregated *de novo* assembly report generated with a custom Python script and 2) evaluation against the FA19 reference genome using QUAST (38, 39). The Python script summarizes standard metrics (e.g., total assembly size, contig count, length-weighted coverage, largest contig, N50/L50, and GC%). This step produces contiguity and accuracy statistics for downstream review.

Following species identification, reference mapping, and assembly statistics generation, a secondary QC filter excludes low-quality and non-*Ng* samples. Samples that fail species, coverage, or assembly quality criteria (Table 1b) are immediately disqualified and do not proceed to subsequent analyses.

#### 4. AMR Subworkflow

The AMR subworkflow of neissflow detects and characterizes resistance-associated genetic features in *Ng* and performs strain typing using a combination of established tools and custom scripts. The steps of the AMR subworkflow as well as the necessary upstream analysis steps can be seen in Figure 2.

**Figure 2:**
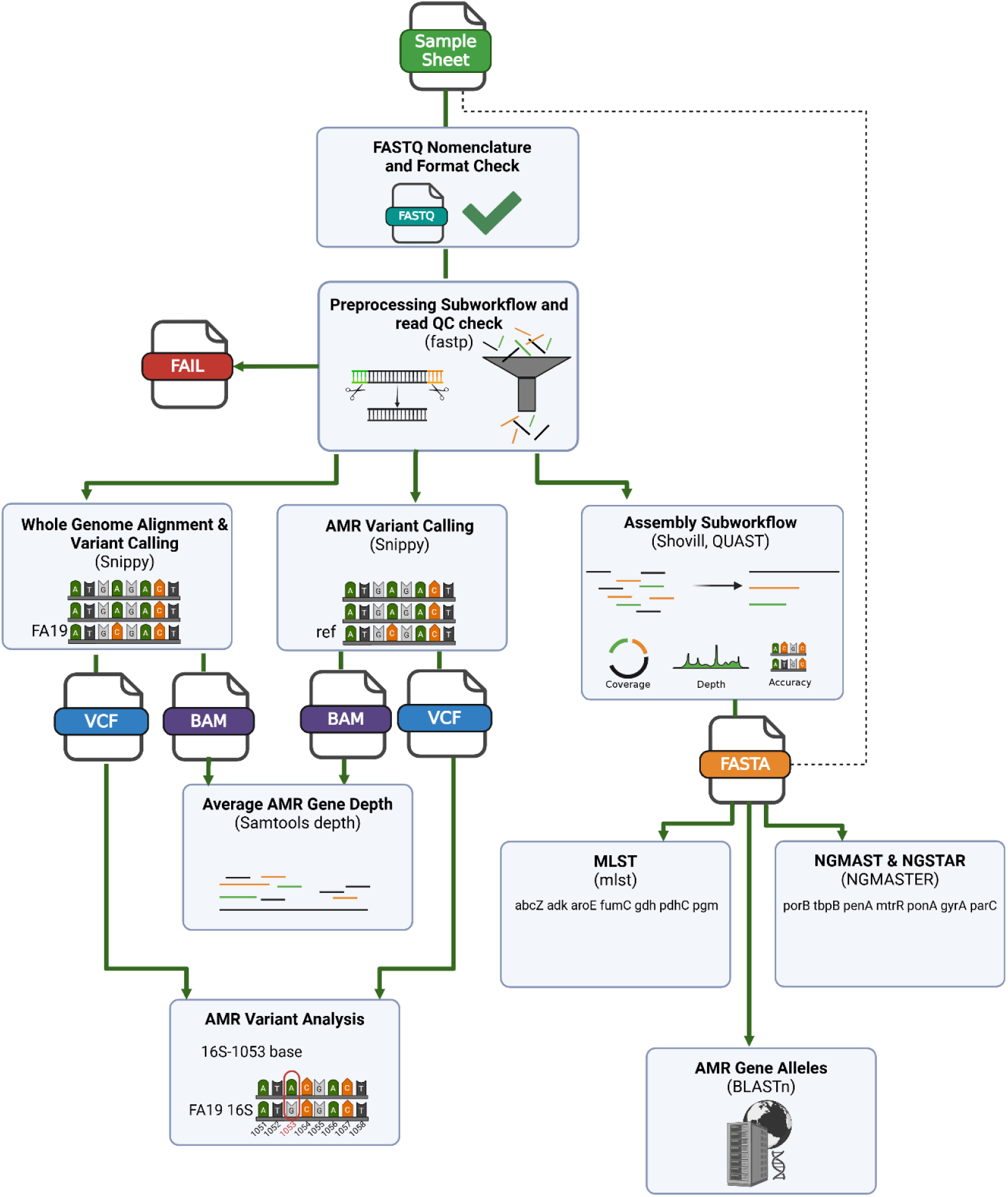
Workflow diagram including the necessary preprocessing and input for the AMR Profiler sub-workflow (assemblies and whole genome alignment and variant calling trimmed and filtered reads) as well as the steps of the AMR Profiler subworkflow (plasmid AMR gene variant calling, AMR gene depth, analysis of AMR variants, Multilocus Sequence Typing (MLST), Ng Multi-Antigen Sequence Typing (NG-MAST), Ng Sequence Typing for Antimicrobial Resistance (NG-STAR), and allele calling). Created with BioRender.com. **QC** — Quality Control; **BAM** — Binary Alignment Map; **VCF** — Variant Call For-mat.

Variant calling is performed using Snippy (v4.6.0) (https://github.com/tseemann/snippy). This step generates variant call format (VCF) files used for downstream AMR gene interrogation. To assess lineage background and strain typing, mlst (v2.23.0) (https://github.com/tseemann/mlst) is employed for multi-locus sequence typing, while NGMASTER (v0.5.6) (40) is used to assign both NG-MAST (multi-antigen sequence typing) and NG-STAR (sequence typing for AMR) profiles. For locus-specific characterization, BLASTn (41) searches are conducted against curated reference databases to identify *penA* and *porB* alleles and to determine the presence of mosaic *mtrR*. The associated databases provided with the repository, and used to perform the validation, were last updated March 30^th^, 2026. Gene and position-specific sequencing depth is assessed with Samtools depth (v1.17) (37), which provides per-position depth of coverage.

A custom Python script integrates outputs from Snippy and Samtools to assess variant calls for AMR-associated loci. The script evaluates a curated panel of chromosomal and plasmid-mediated resistance determinants (Supplementary Table S1). Variant calls are interpreted using depth thresholds and defined criteria for resistance-associated loci (Supplementary Methods).

#### 5. Phylogeny Subworkflow

The phylogeny subworkflow generates phylogenetic trees from a core-genome alignment derived from per-sample variant calls relative to a reference genome (FA19 by default). The steps of the phylogeny sub-workflow can be found in Figure 3. Variant calling and core alignment generation are performed using Snippy (v4.6.0). Sites of recombination are then masked using Gubbins (v3.3.5) (40). Pairwise SNP distances are calculated with snp-dists (v0.8.2) (https://github.com/tseemann/snp-dists), and outbreak clusters are inferred using a graph-based approach implemented using the disjoint set union (DSU) algorithm (Supplementary Methods).

**Figure 3:**
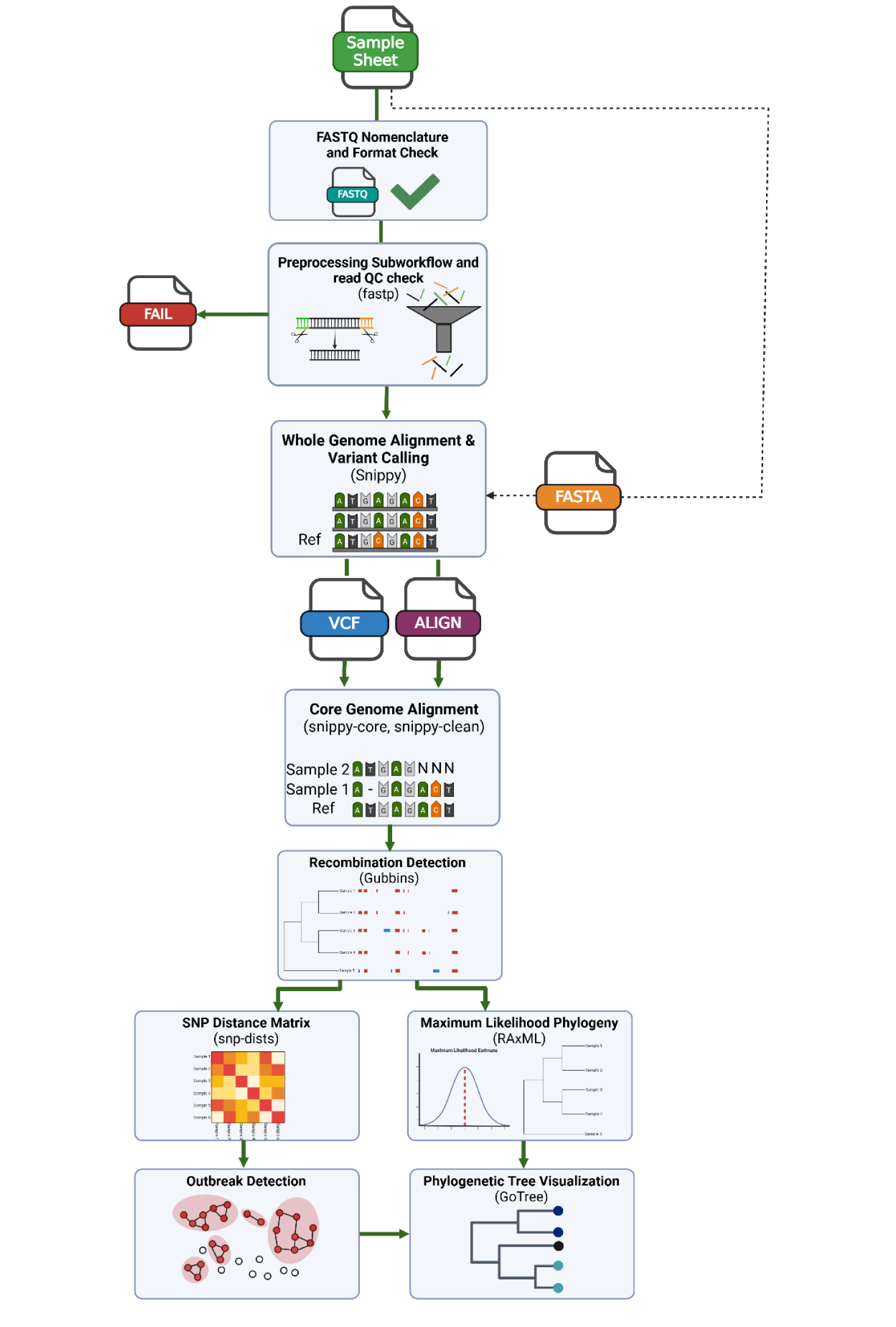
Workflow diagram including the necessary preprocessing steps / input options (trimmed and filtered reads or assemblies) for the Phylogeny subworkflow, as well as the phylogenetic analysis steps (core genome alignment, recombination detection, SNP distance calculation, outbreak detection, maximum likelihood phylogenetic tree generation, and phylogenetic visualization). Created with BioRender.com. **SNP** — Single Nucleotide Polymorphism; **BAM** — Binary Alignment Map; **VCF** — Variant Call Format.

Maximum-likelihood phylogenies are inferred using RAxML-NG (v1.2.2; GTR+Γ with ascertainment bias correction) (40), followed by midpoint rooting with GoTree (v0.4.5) (42). Resulting trees are visualized with cluster-based tip coloring, and summary outputs are provided in an HTML report. Additional implementation details, including alignment processing, ascertainment bias correction, and quality control procedures, are described in the Supplementary Methods.

### Validation design overview

We performed a multi-component validation of neissflow (v2.1.0) using two independent datasets: Dataset 1 evaluated sensitivity and specificity for 10 analytes, and Dataset 2 evaluated reproducibility of all 10 analytes across replicate sequences from 17 *Ng* reference strains. The 10 analytes comprised: 1) *Ng* species identification, 2) MLST, 3) *penA* allele, 4) *porB* allele, 5) 23S rRNA A2059G, 6) 23S rRNA C2611T, 7) *mtrR* promoter mutations, 8) *gyrA* S91 mutations, 9) *blaTEM* plasmid gene, and 10) *tetM* plasmid gene. Ng species identification and MLST represent aggregate analytes. Ng species identification is evaluated as a single integrated call derived from Mash-based genomic screening and read mapping to the FA19 reference genome. MLST is evaluated at the level of the ST assignment, the bioinformatically integrated result of querying all seven housekeeping loci simultaneously. An incorrect allele call at any locus propagates as a discordant ST, ensuring that ST-level concordance is analytically rigorous. Detailed analyte definitions for true positive (TP), true negative (TN), false positive (FP), and false negative (FN) classifications are provided in Supplementary Table S2. Performance was evaluated against prespecified acceptance criteria requiring ≥95% sensitivity, ≥95% specificity, and ≥95% concordance.

### Dataset 1: Analytical Sensitivity and Specificity

#### Dataset Composition

Dataset 1 comprised 158 WGS: 105 *Ng* samples and 53 non-gonococcal *Neisseria* and related species (*N. meningitidis*, *N. lactamica*, *N. subflava*, *N. sicca*, *N. mucosa*, *N. flavescens*, *N. cinerea*, *N. macacae*, *N. polysaccharea*, *N. oralis*, *N. weaveri*, *N. bacilliformis*, *N. elongata*, *N. canis*, *Kingella denitrificans*, *Moraxella catarrhalis*). The 105 *Ng* samples for this study were purposively selected from isolates collected and published under Gonococcal Isolate Surveillance Project (GISP), enhanced GISP (eGISP) and/or Strengthening the United States Response to Resistant Gonorrhea (SURRG) project to ensure analyte representation, with supplemental isolates added for underrepresented 23S rRNA analytes (Supplementary Table S3). Of these 105 Ng isolates, 13 were sourced from the CDC & FDA Antimicrobial Resistance (AR) Isolate Bank *Neisseria gonorrhoeae* Ciprofloxacin Panel (NCIP; Panel ID: 1156; https://wwwn.cdc.gov/ARIsolateBank/Panel/PanelDetail?ID=1156), selected to ensure adequate representation of ciprofloxacin susceptibility profiles and *gyrA*/*parC* gene variants in the validation dataset. Species identification for all 105 Ng isolates was confirmed by MALDI-TOF mass spectrometry prior to WGS. Of the 53 non-Ng and related species isolates, 23 were sourced from the CDC & FDA AR Isolate Bank *Neisseria* Species Panel (Panel ID: 1153; https://wwwn.cdc.gov/ARIsolateBank/panel/paneldetail?ID=1153), from which genome data were generated; genome data for the remaining isolates were obtained from published resources (43). For CDC & FDA AR Isolate Bank isolates, species identification was confirmed by MALDI-TOF mass spectrometry as the primary method; where MALDI-TOF identification was ambiguous or the species was absent from the reference library, confirmation was performed by 16S rRNA gene sequencing and/or whole-genome sequencing-based taxonomic classification, as previously described (42–44). All genome assemblies were submitted to PubMLST for genotype/AMR assignments.

#### Benchmark Standards

Species identification was evaluated against three independent methods: MALDI-TOF mass spectrometry (45), PubMLST ribosomal MLST (46), and PubMLST *rplF* (47) geno-species assignment. AMR genotypes and sequence typing analytes were assessed against PubMLST curated allele and mutation calls from WGS data. For select markers with well-established genotype-phenotype relationships (*gyrA* S91 mutations (ciprofloxacin; (48)), 23S rRNA mutations (azithromycin; (49)), *tetM* (tetracycline; (50)), *blaTEM* (penicillin; (51)), Minimum Inhibitory Concentrations (MICs) values generated either at the Antimicrobial Resistance Laboratory Network (AR Lab Network) and at CDC (52) served as secondary phenotypic validation standards.

#### Performance Metrics

For each analyte, sensitivity [TP/(TP+FN)], specificity [TN/(TN+FP)], positive predictive value [TP/(TP+FP)], negative predictive value [TN/(TN+FN)], and overall concordance [(TP+TN)/total] were calculated with 95% confidence intervals using the Wilson score method. Species identification was assessed using all 158 samples; AMR/typing analytes were evaluated using 105 *Ng* samples.

### Dataset 2: Reproducibility and Precision

#### Dataset Composition

Dataset 2 comprises 283 WGS replicates from 17 *Ng* reference strains. Each strain was sequenced as biological triplicates across multiple laboratories participating in the CDC AR Lab Network participated in the *Ng* WGS External Quality Assessment (EQA) program (2020-2024) (53). In total, each strain was sequenced 15-18 times each (mean=16.6 replicates/strain). The 17 *Ng* reference strains for this study were selected from published international reference isolates (54, 55) (Supplementary Table S4).

### Comparators

PacBio sequences for four reference strains (CDC 10328, F-28, SPL-4, SPJ-15) were generated inhouse, submitted to PubMLST, and used as comparators. For the remaining 13 strains, benchmark genotypes were obtained from peer-reviewed publications reporting PacBio sequence characterization of these reference strains (54, 55).

### Reproducibility Calculation

For each analyte and strain, reproducibility was calculated as the proportion of sequencing replicates producing concordant calls with the benchmark genotype: reproducibility = (number of concordant replicates) / (total replicates). Overall reproducibility was calculated across all strains, all analytes, and all evaluable calls.

### Validation of the Phylogenetic Subworkflow

Of the 283 isolates in Dataset 2, 281 passed QC. Replicates of SPJ-15 (n=15) were excluded, as SPJ-15 is the same strain as NG-P, yielding 266 isolates for validation of the phylogenetic reconstruction and outbreak detection components of neissflow. This final set represented 16 reference strains. Two complementary analyses were performed to assess clustering accuracy at different phylogenetic resolutions.

### Analysis 1: All-Strain Phylogenetic Clustering

All samples were aligned to the FA19 reference genome. The expectation was that all replicates of each strain would form a distinct monophyletic clade with within-clade pairwise SNP distances ≤20 SNPs, consistent with established thresholds for identifying clonally related samples in outbreak investigations (56). Performance was assessed by: 1) correct assignment of all replicates to their respective strain-specific clades, 2) within-clade mean SNP distances ≤20, and 3) no intermixing of replicates from different strains.

### Analysis 2: Intra-MLST Phylogenetic Resolution

To evaluate neissflow’s ability to resolve closely related strains, phylogenetic analyses were performed within three MLST groups using MLST specific references (Supplementary methods)

In intra-MLST analyses, replicates were expected to form distinct monophyletic clusters with pairwise SNP distances ≤12 SNPs (57), consistent with the higher phylogenetic resolution achieved using closely related reference genomes. This lower SNP threshold accounts for the reduced genetic diversity within a single MLST.

For both analyses, phylogenetic accuracy was defined by: 1) correct recovery of all expected strain-specific clusters, 2) confinement of each cluster to SNP distances below the specified threshold (≤20 for inter-strain, ≤12 for intra-MLST), and 3) monophyly of all strain-specific groups with no intermixing of replicates from different strains.

## Results

### Computational Performance

neissflow performed efficiently across datasets of varying sizes under shared CDC HPC resource constraints. Notably, we limited pipeline execution to only 12 parallel jobs, which is common practice in our shared resource environment. Computational resources were allocated to processes based on their assigned process label. The lowest label corresponds to 1 central processing unit (CPU) and 12 GB of memory, and the highest corresponds to 16 CPUs and 128 GB of memory. Using standardized parameters (Gubbins with 5 iterations and RAxML-NG with 500 bootstrap replicates), the pipeline completed the full analysis in 56 minutes 57 seconds for 25 isolates, 7 hours 50 minutes for 158 isolates (Dataset 1; Supplementary Table S3), and 20 hours 44 minutes for 283 isolates (Dataset 2; Supplementary Table S4). neissflow has also been successfully run end to end on a Windows laptop (Dell Inc. XPS 15 9520) using Windows Subsystem for Linux (WSL) with 14 Cores (12^th^ Gen Intel® Core™ i9-12900HK, 2500 Mhz) and 64.0 GB of installed physical memory (RAM) with a 7 sample test set in 10 minutes and 29 seconds, with FASTQ and FASTA input.

### Species Identification Performance

neissflow demonstrated accurate *Ng* species identification across 158 isolates, comprising 105 *Ng* and 53 non-gonococcal *Neisseria* and related species (Table 2; Supplementary Tables S5 and S3). When evaluated against three independent methods (MALDI-TOF mass spectrometry, PubMLST ribosomal MLST typing, and PubMLST *rplF* genospecies assignment) neissflow achieved 100% sensitivity (95% CI: 96.6-100%), 100% specificity (95% CI: 93.3-100%), and 100% overall concordance (95% CI: 97.7-100%) for *Ng* identification, corresponding to a percent positive agreement (PPA) of 100% and percent negative agreement (PNA) of 100% against all three reference methods (Table 2). One *Ng* isolate (GCWGS-21761) failed to receive a species assignment from ribosomal MLST due to one undetermined locus (BACT00003/53); however, this isolate was correctly identified as *Ng* by neissflow and confirmed by both MALDI-TOF and *rplF* genospecies assignment.

**Table 2.**
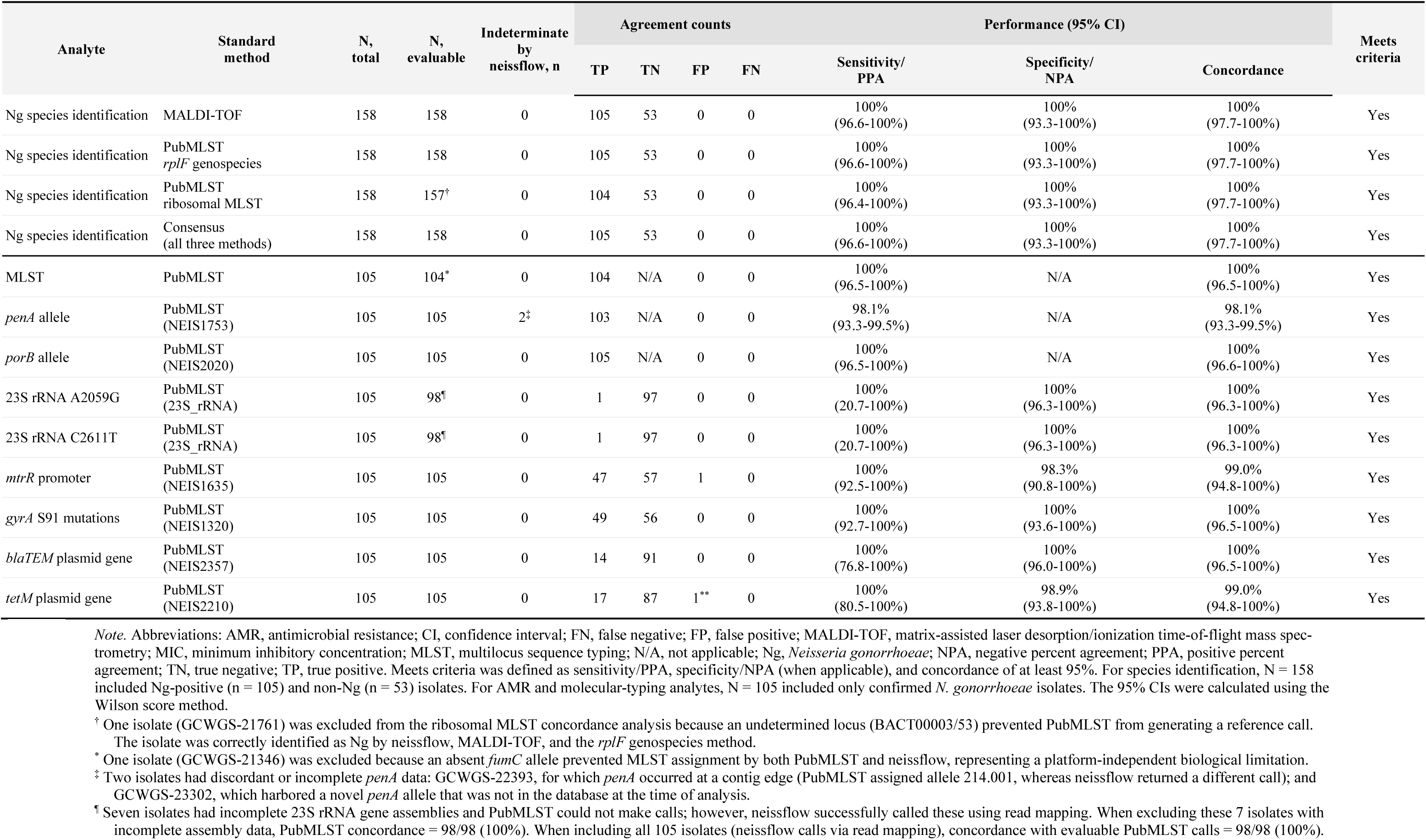
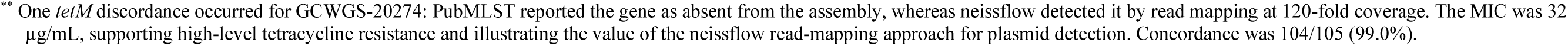
Concordance Between neissflow-Detected 10 analytes and Comparator Methods.

## AMR Marker and Molecular Typing Performance

### Overview of Genotypic Validation

Nine AMR and molecular typing analytes were validated using 105 *Ng* isolates from Dataset 1, with PubMLST genotypic assignments serving as the primary standard (Supplementary Table S2). All 9 analytes achieved ≥95% overall concordance with PubMLST, meeting prespecified acceptance criteria. Seven of 9 analytes demonstrated 100% concordance (Table 2, Supplementary Tables S6 and S3). Three analytes showed minor discordances: *penA* allele (98.1% concordance, 103/105), *mtrR* promoter mutations (99.0% concordance, 104/105) and *tetM* plasmid gene detection (99.0% concordance, 104/105).

### Molecular Typing Markers

#### MLST

neissflow achieved 100% concordance (104/104 evaluable isolates) with PubMLST for multi-locus sequence typing. One isolate (GCWGS-21346) lacked a *fumC* allele call in both PubMLST and neissflow, preventing MLST assignment in both platforms. All successfully typed isolates showed perfect agreement between neissflow and PubMLST sequence type assignments.

#### *penA* allele

neissflow demonstrated 98.1% concordance (103/105) for *penA* allele assignment. Two isolates required special consideration: GCWGS-22393 had the *penA* gene at a contig edge, resulting in discordant allele calls between PubMLST (214.001) and neissflow (358.001); while neissflow assigned *penA* allele 2.001 for GCWGS-23302 but PubMLST did not assign a *penA* allele at the time of analysis. Excluding these two technical challenges, concordance was perfect.

#### *porB* allele

neissflow achieved 100% concordance (105/105) with PubMLST for *porB* allele typing. All 105 isolates showed perfect agreement between neissflow and PubMLST *porB* allele assignments.

### Chromosomal AMR Markers

#### 23S rRNA mutations (A2059G and C2611T)

Both 23S rRNA mutations associated with azithromycin resistance showed 100% concordance with PubMLST among isolates with complete 23S rRNA gene assemblies (98/98 isolates). Seven isolates had incomplete 23S rRNA genes in their assemblies, preventing PubMLST from making calls. However, neissflow successfully detected wild type 23S rRNA sequences in all seven cases using read mapping to the FA19 reference genome. One isolate (GCWGS-23949) carried A2059G, and one isolate (GCWGS-22115) carried C2611T; both were correctly identified by neissflow and confirmed by PubMLST, with corresponding high azithromycin MICs (16 µg/mL for both).

#### *mtrR* promoter mutations

neissflow achieved 99.0% concordance (104/105) with PubMLST for detection of *mtrR* promoter alterations. Among the 105 isolates, 47 carried promoter mutations and 57 had wild-type promoter sequences.

#### *gyrA* S91 mutations

Perfect concordance (100%, 105/105) was observed between neissflow and PubMLST for *gyrA* amino acid 91 mutation detection. The dataset included 48 isolates with the S91F mutation and one with S91I, both associated with not susceptible ciprofloxacin MICs (0.125-64 µg/mL). It also included 51 isolates with wild-type S91 and five with the S91T mutation. S91T was classified as wild type because it is associated with susceptible CIP MIC (0.001-0.060 µg/mL). All nucleotide calls were concordant across methods. Notably, PubMLST did not report the S91I mutation for isolate GCWGS-23949 in its summary output, although the mutation was present in the *gyrA* sequence annotated by PubMLST; neissflow correctly identified this mutation.

### Plasmid-Mediated AMR Markers

#### *blaTEM* plasmid

neissflow demonstrated 100% concordance (105/105) with PubMLST for *blaTEM* β-lactamase gene detection. Fourteen isolates carried the *blaTEM* gene, and 91 isolates lacked the gene; all calls were concordant between platforms.

#### *tetM* plasmid

Overall concordance was 99.0% (104/105) with one notable discordance. Seventeen isolates carried *tetM* gene in their plasmid conferring tetracycline resistance, and 87 lacked the gene. One isolate (GCWGS-20274) was *tetM*-positive by neissflow using read mapping (120x coverage depth) but *tetM*-negative by PubMLST from the assembly. Phenotypic data strongly supported neissflow’s call: this isolate showed high-level tetracycline resistance (MIC = 32 µg/mL), consistent with plasmid-mediated resistance.

### Genotype-Phenotype Correlations

To assess the clinical relevance of neissflow AMR marker detection, we evaluated concordance between genotypic markers and antimicrobial susceptibility phenotypes (MIC values) for four antimicrobials (Supplementary Tables S6 and S7). One isolate, GCWGS-20890, lacked MIC data and was excluded from these analyses.

#### Ciprofloxacin

##### *gyrA* S91F and S91I Mutations

Detection of ciprofloxacin resistance associated *gyrA* S91 mutations (S91F and S91I) by *neissflow* showed complete concordance with either intermediate or resistant phenotypes (MIC ≥0.125 µg/mL), with 100% sensitivity (95% CI: 92.6-100%), 100% specificity (95% CI: 93.6-100%), and 100% overall agreement (95% CI: 96.5-100%). Among 56 isolates without resistance associated mutations (S91 wild-type, n=51; S91T, n=5), 56 (100%) had MIC <0.06 µg/mL (range: 0.001–0.03 µg/mL), indicating susceptibility (Supplementary Table S7).

#### Azithromycin

##### 23S rRNA Mutations

Detection of 23S rRNA mutations (A2059G or C2611T) showed perfect genotype-phenotype correlation (100% sensitivity, 100% specificity, 100% concordance) when using MIC ≥16 µg/mL as the breakpoint for high-level azithromycin resistance. Both isolates carrying 23S mutations (GCWGS-23949 with A2059G; GCWGS-22115 with C2611T) exhibited high-level resistance (MIC = 16 µg/mL). All 102 isolates without 23S rRNA mutations had MIC ≤ 2 µg/mL, with zero false positives or false negatives (Supplementary Tables S6 and S7).

#### Tetracycline

##### *tetM* Plasmid

Detection of the *tetM* gene yielded perfect genotype-phenotype correlation (100% sensitivity, 100% specificity, 100% concordance) when using MIC ≥16 µg/mL as the cuttoff for plasmid-mediated high-level tetracycline resistance. All 17 *tetM*-positive isolates exhibited high-level tetracycline resistance (MIC range: 16-64 µg/mL, median 32 µg/mL), suggesting plasmid-mediated resistance. All 87 *tetM*-negative isolates had MIC <16 µg/mL (Supplementary Tables S6 and S7).

#### Penicillin

##### *blaTEM* Plasmid

*blaTEM* gene detection by neissflow showed high specificity (96.3%; 95% CI: 89.4-99.2%) but moderate sensitivity (45.8%; 95% CI: 25.6-67.2%) for predicting penicillin MIC ≥2.0 µg/mL, achieving 84.6% overall concordance (95% CI: 76.4-90.9%). Among 14 *blaTEM*-positive isolates detected by neissflow, 10 (71.4%) showed elevated penicillin MICs (range: 8-64 µg/mL), confirming β-lactamase-mediated resistance. Four *blaTEM*-positive isolates had MICs <2.0 µg/mL (range: 0.19-1.0 µg/mL). Thirteen isolates had penicillin MIC ≥2.0 µg/mL (range: 2.0-4.0 µg/mL) without detectable *blaTEM* (Supplementary Tables S6 and S7).

### Reproducibility and Precision: Dataset 2

Dataset 2 comprised 283 whole-genome sequences from 17 *Ng* reference strains sequenced 15-18 times each across multiple AR Lab Network *Ng* Regional Laboratories. Of 283 sequences, 281 (99.3%) passed neissflow’s two-stage QC process. Two sequences (GCWGS-28790 from NG-M; GCWGS-15877 from NG-X) failed Stage 2 QC due to assembly quality metrics. Species identification succeeded for all 283 sequences (100%), including the two QC failures.

#### Per-Analyte Reproducibility

Nine of ten analytes achieved 100% reproducibility with expected results across all evaluable replicates (Table 3): species identification (283/283), MLST (281/281), *porB* allele (281/281), 23S rRNA mutations (281/281), *mtrR* promoter (281/281), *gyrA* S91 (281/281), *blaTEM* gene (281/281), and *tetM* gene (281/281). *penA* allele typing showed 99.6% reproducibility (280/281), with one discordant call in strain NG-L replicate GCWGS-29616 (assigned allele 17.001 instead of expected 7.001) (Supplementary Table S8).

**Table 3.** Summary of Reproducibility Performance of neissflow by Analyte (Dataset 2)

| Analyte | Total Repli-<br>cates | Concord-<br>ant Calls | Discordant<br>Calls | QC Fail-<br>ures* | Reproducibil-<br>ity (%) | 95%<br>CI** |
| --- | --- | --- | --- | --- | --- | --- |
| Species ID (*Ng*) | 283 | 283 | 0 | 0 | 100% | 98.7-100% |
| MLST | 281 | 281 | 0 | 2 | 100% | 98.7-100% |
| *penA* allele | 281 | 280 | 1 | 2 | 99.6% | 98.0-100% |
| *porB* allele | 281 | 281 | 0 | 2 | 100% | 98.7-100% |
| 23S rRNA A2059G | 281 | 281 | 0 | 2 | 100% | 98.7-100% |
| 23S rRNA C2611T | 281 | 281 | 0 | 2 | 100% | 98.7-100% |
| *mtrR* promoter | 281 | 281 | 0 | 2 | 100% | 98.7-100% |
| *gyrA* S91 | 281 | 281 | 0 | 2 | 100% | 98.7-100% |
| *blaTEM* plas-<br>mid gene | 281 | 281 | 0 | 2 | 100% | 98.7-100% |
| *tetM* plasmid gene | 281 | 281 | 0 | 2 | 100% | 98.7-100% |
| **Overall (all ana-**<br>**lytes)** | **3,093\*** | **3,092\*** | **1\*** | **20\*** | **99.97%** | **99.9-**<br>**100%** |

\* QC failures represent 2 replicates (one from NG-M, one from NG-X) that failed Stage 2 QC.
\*\* 95% confidence intervals calculated using the Wilson score method.

#### Per-Strain Reproducibility

Sixteen of 17 strains (94.1%) demonstrated 100% reproducibility across all replicates and analytes (Supplementary Table S8). One strain, NG-L, showed <100% reproducibility with one *penA* discordance. For NG-M and NG-X, all replicates passing QC showed 100% concordant analyte calls.

#### Overall Reproducibility

Across all 17 strains, 10 analytes, and 3,093 evaluable calls, neissflow achieved 99.97% overall reproducibility (3,092/3,093 concordant calls; 95% CI: 99.9-100%), exceeding the prespecified ≥95% acceptance threshold for all analytes (Tables 3; Supplementary Table S4).

### Phylogenetic Subworkflow Validation

#### All-Strain Phylogenetic Clustering (Analysis 1)

Phylogenetic analysis of the 266 isolates representing 16 strains yielded monophyletic clades with all replicates correctly grouped and no intermixing between strains (Table 4a, Figure 4). Strains sharing the same MLST formed separate, distinct monophyletic clades. Mean within-clade pairwise SNP distances ranged from 2.9 (NG-P) to 13.63 (F-28), with all strains showing mean distances ≤14 SNPs. Maximum pairwise distances within clades ranged from 7 (NG-P) to 23 SNPs (NG-Z); notably, the two strains exceeding 20 SNPs (NG-Z and F-28) maintained mean SNP differences of 6.96 and 13.63 SNPs, respectively - well within the outbreak threshold and representing a single outlier pairwise comparison.

**Figure 4.**
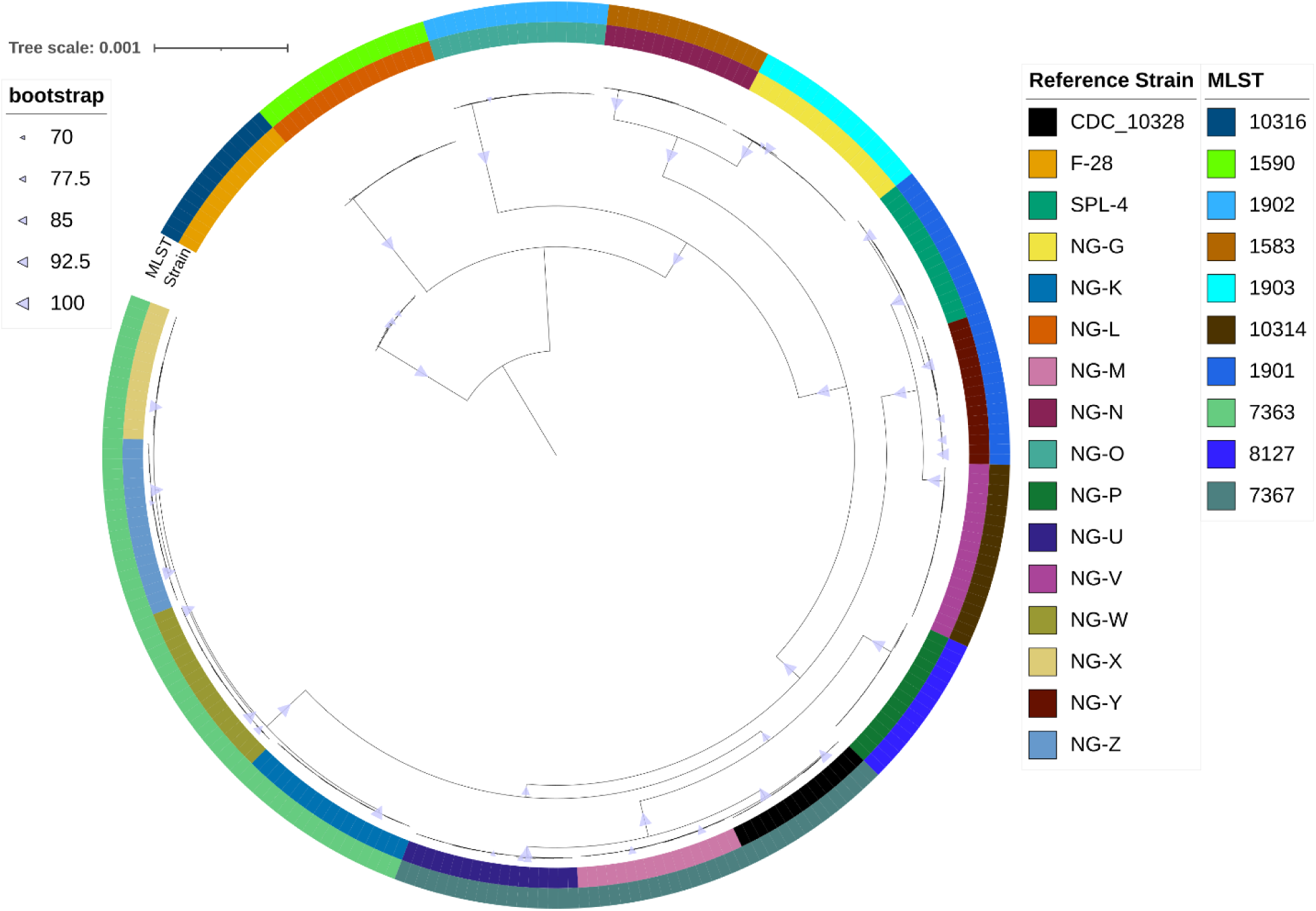
Phylogenetic validation of neissflow using replicate sequences from 16 *N. gonorrhoeae* refer-ence strains. (a) All-strain phylogenetic clustering of 266 isolates representing 16 reference strains, aligned to the FA19 reference genome. Maximum-likelihood phylogenetic tree shows strain-specific clustering with tips colored by reference strain identity (see legend). All 16 strains (100%) formed monophyletic clades with within-strain mean pairwise SNP distances ranging from 3.31 to 13.65 SNPs, all meeting the ≤20 SNP outbreak threshold (see Table 4a for detailed SNP metrics). Scale bar indicates substitutions per site. Tree was constructed using RaxML-NG with GTR+Γ model and ascertainment bias correction, following re-combination removal by Gubbins. Tree is midrooted. Figure generated with iTOL (https://itol.embl.de/, v7.5.1) (69).

**Table 4.** Phylogenetic Clustering Performance: All-Strain and Intra-MLST Analyses.

| Strain/Clade (MLST) | N | Mean<br>SNP | Min<br>SNP | Max<br>SNP | Monophyletic | Meets Crite-<br>ria* |
| --- | --- | --- | --- | --- | --- | --- |
| NG-W (ST-7363) | 18 | 3.11 | 0 | 7 | Yes | Yes |
| NG-X (ST-7363) | 14 | 3.97 | 1 | 8 | Yes | Yes |
| NG-P (ST-8127) ‡ | 15 | 2.9 | 0 | 7 | Yes | Yes |
| NG-N (ST-1583) | 16 | 3.5 | 0 | 9 | Yes | Yes |
| SPL-4 (ST-1901) | 15 | 4.36 | 1 | 9 | Yes | Yes |
| NG-K (ST-7363) | 18 | 4.47 | 0 | 11 | Yes | Yes |
| NG-Y (ST-1901) | 15 | 5.09 | 0 | 12 | Yes | Yes |
| NG-V (ST-10314) | 18 | 4.33 | 1 | 9 | Yes | Yes |
| NG-G (ST-1903) | 18 | 4.68 | 0 | 11 | Yes | Yes |
| NG-O (ST-1902) | 18 | 4.65 | 0 | 10 | Yes | Yes |
| CDC 10328 (ST-7367) | 15 | 6.21 | 0 | 13 | Yes | Yes |
| NG-M (ST-7367) | 17 | 6.61 | 0 | 12 | Yes | Yes |
| NG-U (ST-7367) | 18 | 4.31 | 0 | 10 | Yes | Yes |
| NG-Z (ST-7363) | 18 | 6.96 | 0 | 23 | Yes | Yes** |
| NG-L (ST-1590) | 18 | 7.38 | 0 | 17 | Yes | Yes |
| F-28 (ST-10316) | 15 | 13.63 | 2 | 22 | Yes | Yes** |
| **Overall** | **266** | **4.89** | **0** | **23** | **16/16 (100%)** | **16/16 (100%)** |

\* Criteria: 1) monophyletic clustering, 2) mean within-clade SNP $\leq 20$ , 3) no intermixing.
\*\* NG-Z and F-28 had maximum pairwise distance of 23 and 22 SNPs, respectively but mean SNP distances $\leq 20$ SNPs, meeting mean threshold criterion.
‡ NG-P and SPJ-15 were the same strain and only NG-P were included in this phylogenetic analysis

b. Within-Strain Pairwise SNP Distances for Intra-MLST Analysis Using MLST-Specific References (Analysis 2, 3 MLSTs, n=147, threshold: $\leq 12$ SNPs)
| Strain/Clade (MLST) | N | Mean SNP | Min SNP | Max SNP | Monophyletic | Meets Criteria* |
| --- | --- | --- | --- | --- | --- | --- |
| NG-Y (ST-1901) | 15 | 0.02 | 0 | 1 | Yes | Yes |
| SPL-4 (ST-1901) | 15 | 0.50 | 0 | 2 | Yes | Yes |
| NG-K (ST-7363) | 17 | 0.00 | 0 | 0 | Yes | Yes |
| NG-Z (ST-7363) | 18 | 0.69 | 0 | 5 | Yes | Yes |
| NG-X (ST-7363) | 14 | 0.88 | 0 | 2 | Yes | Yes |
| NG-W (ST-7363) | 18 | 0.99 | 0 | 3 | Yes | Yes |
| NG-U (ST-7367) | 18 | 0.13 | 0 | 2 | Yes | Yes |
| CDC 10328 (ST-7367) | 17 | 0.36 | 0 | 2 | Yes | Yes |
| NG-M (ST-7367) | 17 | 0.24 | 0 | 2 | Yes | Yes |
| **Overall** | **147** | **0.43** | **0** | **5** | **9/9 (100%)** | **9/9 (100%)** |

\* Criteria: 1) monophyletic clustering, 2) mean within-clade SNP $\leq 12$ , 3) no intermixing.

#### Intra-MLST Phylogenetic Resolution (Analysis 2)

Phylogenetic analyses were performed separately for three MLSTs (147 isolates representing nine strains), with each analysis yielding monophyletic clades with low within-cluster SNP diversity. Mean pair-wise SNP distances ranged from 0.00 (NG-K, all replicates identical) to 0.99 (NG-W), with all strains showing mean <1 SNP (Table 4b; Supplementary figure 1). Maximum pairwise distances ranged from 0 (NG-K) to 5 SNPs (NG-Z), well below the prespecified ≤12 SNP threshold.

## Discussion

The results indicate that neissflow provides accurate and reproducible genomic analysis of *Ng* sequences across multiple use-cases relevant to public health surveillance. Validation across two independent datasets showed ≥99% concordance for all benchmark analytes and 100% accuracy for phylogenetic clustering. The outbreak detection algorithm also correctly assigned 100% of isolates to expected strain groups. This level of performance is particularly important in laboratory settings where bioinformatics capacity is limited, and accurate pathogen characterization is critical for public health response. By integrating species confirmation, AMR profiling, molecular typing, and phylogenetic analysis into a workflow with a single-entry point, neissflow helps address practical limitations of existing approaches.

These advantages are particularly evident in AMR detection, where read-based approaches can provide important benefits over *de novo* assembly-based methods. Across key resistance markers, neissflow showed complete genotype-phenotype concordance for well-established associations. For plasmid-borne resistance genes, *de novo* assemblies may incompletely represent plasmid sequences, leading to false-negative calls. This was illustrated by isolate GCWGS-20274, which was *tetM*-positive by neissflow using read mapping (120X depth of coverage), but *tetM*-negative by PubMLST based on the assembly. Phenotypic data supported the neissflow result, with this isolate showing high-level tetracycline resistance (MIC = 32 µg/mL), consistent with plasmid mediated resistance. (Supplementary Tables S7 and S6).

Similar discrepancies were observed for chromosomal AMR markers. For isolate GCWGS-23949, PubMLST did not report the *gyrA* S91I mutation in its summary output, despite the mutation being present in the annotated sequence, whereas neissflow correctly identified it. The corresponding ciprofloxacin MIC indicated resistance, supporting neissflow’s result. This highlights a practical limitation of relying solely on PubMLST summary outputs, which may not surface all or new resistance markers. Assembly-based approaches present additional challenges. Seven isolates had incomplete 23S rRNA gene assemblies, preventing PubMLST from making calls. In contrast, neissflow identified wild type 23S rRNA sequences in all seven cases using read mapping to the FA19 reference genome, avoiding unresolved azithromycin resistance that could impact surveillance efforts. Additionally, neissflow results include the proportion of the four chromosomal 23S rRNA copies carrying A2059G and C2611T mutations. Given the relationship between mutation copy number and azithromycin resistance (49, 58), this information may improve interpretation of resistance phenotypes and surveillance of emerging resistance. Mutation copy number cannot be obtained reliably from assembly-based methods. A further advantage of read-based variant calling is the availability of site-specific quality information. For isolate GCWGS-22115, PubMLST identified *mtrR* allele 39 (ACAAG), whereas neissflow reported NFCAAA due to insufficient coverage (<10X at the first position). This low-confidence call prompted manual inspection in Interactive Genomic Viewer (IGV), which revealed an unreported deletion at that position (true allele: delACAA) (Supplementary Figure 2) (59). Notably, this represents a complex edge case in which neither assembly-based methods nor the current implementation of neissflow fully resolved the correct allele automatically. However, neissflow’s reporting flagged the ambiguity, enabling further investigation and accurate interpretation, whereas the assembly-based result appeared as a definitive but incorrect call.

The AMR validation results also provided insight into the relationship between *blaTEM* plasmid gene carriage and penicillin resistance. Additional analysis revealed a gene-dosage effect, wherein *blaTEM* copy number monotonically increases with penicillin MIC, consistent with existing evidence (60, 61). This relationship may explain the discordances found between *blaTEM*-positive isolates and sensitive phenotypes (Supplementary Figure 3), where lower copy numbers may produce insufficient β-lactamase activity to confer a resistant MIC. Conversely, phenotypically resistant *blaTEM-negative* isolates were also observed, suggesting that penicillin resistance in these samples are likely mediated by chromosomal mechanisms, including *penA* mosaic alleles, *mtrR* mutations, and *ponA* L421P (62, 63) (Table 3, Supplementary Table S4). These findings demonstrate that neissflow reliably identifies resistance-associated genetic markers with sufficient accuracy to support genomic surveillance and AMR trend monitoring in *Ng* populations.

Validation of 23S rRNA A2059G and C2611T mutation calling in Dataset 1 yielded 100% sensitivity and specificity for both analytes (TP=1, TN=97 each); however, we acknowledge that the single true positive per analyte is a shortcoming of the isolate selection in Dataset 1. To address this, reproducibility was assessed in Dataset 2, where strains NG-U (C2611T) and NG-V (A2059G) were each sequenced 18 times and both mutations were accurately called across all replicates (100% reproducibility, 18/18). Further confidence is drawn from the application of neissflow v2.1.0 to 4,084 *Ng* isolates from the US national surveillance program (2023–2025) (manuscript in preparation), in which 3 isolates carrying A2059G and 25 carrying C2611T were detected, with elevated azithromycin, confirming concordance between genotypic and phenotypic resistance (Supplementary Table S9).

Although neissflow reports 81 AMR determinants and sequence typing outputs, formal validation was performed on 10 representative analytes selected based on clinical priority and computational methodology. These analytes encompass the major resistance mechanisms relevant to gonorrhea treatment with well-established genotype-phenotype correlations with MIC elevations, distinguishing them from the remaining determinants which largely represent putative resistance associations. Methodologically, all 81 outputs derive from either genome assembly-based allele calling or read mapping-based variant detection, and the 10 representative analytes were selected to validate both pathways, providing workflow-level validation (64–66).

The phylogenetic validation assessment further demonstrated the utility of neissflow. Across replicate datasets, the pipeline consistently recovered expected strain level clustering, with accurate reconstruction of monophyletic clades. Evaluation of intra-MLST datasets showed that neissflow resolves closely related isolates with low SNP diversity, supporting its ability to detect fine scale genomic differences. The use of MLST-specific reference genomes rather than the distant FA19 reference maximized phylogenetic resolution, enabling clear discrimination of strains, underscoring the importance of reference selection in outbreak investigations.

While several pipelines exist for bacterial phylogenetic analysis (e.g., bactmap (https://github.com/nf-core/bactmap), UBCG2 (67), PhyloPhlAn (68)), these pipelines are species agnostic and require configuration by the user to perform *Ng* analysis. Additionally, they would need to be used in concert with other tools to gain the contextual information output by neissflow (AMR, ST, predicted outbreak clustering, etc.). By default, the pipeline compares genomes to the *Ng* reference genome FA19, alleviating the need to provide a reference unless desired. The outbreak detection analysis also provides interpretable clusters alongside the tree to aid epidemiological inference for surveillance. However, genomic similarity does not necessarily imply epidemiological linkage; neissflow’s graph-based algorithm flags networks of closely related isolates, but outbreak confirmation requires epidemiological context.

In addition to analytical performance, neissflow is designed for accessibility and ease of use in public health settings. It requires minimal input preparation (sample sheet with FASTQ file paths and basic parameters), handling all downstream processing, quality control, and result generation. Comprehensive HTML reports with visualizations (phylogenetic trees with color-coded outbreak clusters, SNP distances, QC metrics) enable epidemiologists without bioinformatics training to interpret results. The nf-core framework ensures best-practice pipeline structure, with standardized formats, documentation, and dependency management, while supporting execution across local, HPC, and cloud environments. This flexibility allows resource limited laboratories to leverage scalable cloud infrastructure (e.g., AWS, Google Cloud, Azure) without local HPC investment, expanding access to genomic surveillance.

Taken together, neissflow addresses key limitations of existing *Ng* genomic analysis tools by providing a fully integrated, automated pipeline that performs species identification, chromosomal and plasmid AMR detection, molecular typing, and phylogenetic analysis from raw sequencing reads. This end-to-end, single entry point workflow eliminates the need to combine multiple tools (PubMLST for typing, separate phylogenetic software, manual AMR annotation) or possess tool specific expertise, barriers that have limited genomic surveillance implementation in resource-constrained public health laboratories.

neissflow currently only supports short-read Illumina sequencing data. In the future, we plan to support long-read sequencing inputs (e.g. Oxford Nanopore, PacBio) with the aim of adding the ability to generate more complete draft genome assemblies and supporting PHLs with these sequencing data types. Furthermore, variant calling relies strictly on the FA19 reference genome, which does not capture the total breadth of gene content diversity of *Ng* (e.g. plasmids) and may reduce sensitivity at highly divergent loci. Similarly, AMR detection depends on select, well-characterized resistance determinants, which will not capture the gonococcal resistome in its entirety. neissflow’s AMR detection also requires ≥10x coverage at each position for confident variant calling which may reduce sensitivity for low-coverage samples. Future enhancements include integration of machine learning for novel AMR marker discovery, supporting long-read sequencing data, and direct Laboratory Information Management System (LIMS) integration for seamless laboratory workflows. Application Programming Interface (API) development would enable automated data exchange with epidemiological databases (Combating Antimicrobial Resistant Gonorrhea and other STIs (CARGOS; https://www.cdc.gov/sti/php/projects/cargos.html), Pub-MLST), facilitating real-time national and global surveillance.

## Conclusion

neissflow provides the *Ng* research and public health communities with a robust and scalable end-to-end bioinformatics pipeline for comprehensive genomic analysis. Exceptional analytical performance (≥99% concordance, 99.97% reproducibility, 100% phylogenetic accuracy) and user-friendly design position neissflow as a reference standard for *Ng* genomic surveillance. Open-source availability encourages community adoption, collaborative development, and continuous improvement. Our findings demonstrate the utility of neissflow as a reproducible and scalable workflow for genomic characterization of *Ng* and AMR-associated determinants, supporting its potential application in genomic surveillance and public health laboratories with appropriate validation and implementation considerations.

## Acknowledgements

We thank the Gonococcal Isolate Surveillance Project (GISP), Enhanced GISP, and the Strengthening the US Response to Resistant Gonorrhea (SURRG) program for contributing isolates used in our analyses. We thank the 4 AR Laboratory Network regional laboratories (https://www.cdc.gov/antimicrobial-resistance-laboratory-networks/php/about/index.html); Washington State Department of Health, WA; Maryland Department of Health, MD; Tennessee Department of Health, TN; and Utah Public Health Laboratory, UT for generating WGS and AST data for the isolates included in this study. We thank Alesia Harvey for her assistance in verifying isolate epidemiological and minimum inhibitory concentration (MIC) data. We thank Brian Raphael and Ellen Kersh for their support in funding acquisition and for their guidance during the initial stages of this project. We thank Hsi Liu and Cau D. Pham for sequencing some of the *Neisseria commensal* isolates from the CDC & FDA Antimicrobial Resistance Isolate Bank; *Neisseria* Species Panel (Panel ID: 1153) used in this analysis. We thank Amantha Smith for her contributions to the initial figures presented in this manuscript. We thank the Office of Advanced Molecular Detection’s Scientific Computing Team for their technical support.

## Ethical statement

The GISP, eGISP and SURRG protocols and projects are reviewed by the Office of the Associate Director for Science (ADS), NCHHSTP, CDC periodically. Most recently, this was done in February 2021 (for GISP, eGISP) and 2023 (for SURRG). The GISP, eGISP and SURRG projects were determined to be surveillance activities and not human subjects research.

## Disclaimer

The findings and conclusions in this report are those of the authors and do not necessarily represent the official position of the Centers for Disease Control and Prevention.

## Data Availability

Documentation for the usage of neissflow, including explanations of all features, is available at https://github.com/CDCgov/neissflow. The validation datasets are publicly available for laboratories to perform inhouse validation of neissflow implementation (NCBI SRA BioProject PRJNA1367095; https://www.ncbi.nlm.nih.gov/bioproject/PRJNA1367095). Supplementary Tables S5 and S6 lists the individual accessions for the whole genome data for all the isolates with associated metadata used in this study. Some of the isolates used in this study are available at the CDC & FDA Antimicrobial Resistance Isolate Bank: (1) the *N. gonorrhoeae* Ciprofloxacin Panel (NCIP; Panel ID: 1156; https://wwwn.cdc.gov/ARIsolateBank/Panel/PanelDetail?ID=1156), comprising 14 *N. gonorrhoeae* isolates representing a wide range of ciprofloxacin susceptibility profiles and *gyrA/parC* gene variants, used to supplement the validation dataset for fluoroquinolone resistance markers; and (2) the *Neisseria* Species Panel (Panel ID: 1153; https://wwwn.cdc.gov/ARIsolateBank/panel/paneldetail?ID=1153), comprising representative *Neisseria* species including *N. gonorrhoeae*, *N. meningitidis*, and other *Neisseria* commensals.

## Supplementary Figures

**Supplementary Figure 1:** Phylogenetic validation of neissflow using replicate sequences. (A) Intra-MLST phylogenetic clustering for MLST 1901 (n=30 isolates) from two reference strains (NG-Y, SPL-4), aligned to the NG-Y reference genome. Both strains formed distinct monophyletic clades with 100% accuracy; mean within-strain SNP distance was 0.26, with all pairwise comparisons ≤2 SNPs (see Table 4b). (B) Intra-MLST phylogenetic clustering for MLST 7363 (n=67 isolates) from four reference strains (NG-K, NG-W, NG-X, NG-Z), aligned to the NG-K reference genome. All four strains formed distinct monophyletic clades with 100% accuracy; mean within-strain SNP distance was 0.43, with all pairwise comparisons ≤5 SNPs (see Table 4b). (C) Intra-MLST phylogenetic clustering for MLST 7367 (n=50 isolates) from three reference strains (CDC 10328, NG-M, NG-U), aligned to the NG-U reference genome. All three strains formed distinct monophyletic clades with 100% accuracy; mean within-strain SNP distance was 0.24, with all pairwise comparisons ≤2 SNPs (see Table 4b). Tips are colored by reference strain identity (see legends). All trees were constructed using RAxML-NG with GTR+Γ model and ascertainment bias correction, following recombination removal by Gubbins. Trees are midpoint rooted. The use of MLST-specific reference genomes maximizes phylogenetic resolution and enhances detection of recent recombination events compared to the distantly related FA19 reference used in Figure 4. Figure generated using iTOL (v7.5.1) (62) and Inkscape.

**Supplementary Figure 2:** GCWGS-22115 alignment to FA19 showing the *mtrR* promoter (1110844-1110848) visualized in IGV. This was reported as NFCAAA by neissflow and ACAAG by PubMLST. This deletion would cause the true *mtrR* allele to be delACAA. neissflow parses Snippy output to make this determination. The deletion was reported by Snippy as a complex variant at the second position, resulting in a change from AA to C. In other cases where a deletion has been identified at this position, Snippy has reported a deletion at the position before the start of the *mtrR* promoter sequence (TA to T), and thus an explicitly indicated deletion is what neissflow looks for to report a “del”. Even if neissflow had correctly parsed this edge case, the reported variant would have been ACdelAA, since Snippy reports complex variants at the position that they start. This also appears to be inaccurate based on what can be seen at these positions with IGV. PubMLST reports the allele as ACAAG as that is what resolves into the assembly. The G that they are reporting in the last position of the allele is the G from one position over and is not a true SNP (wild type is AAAAA flanked by a T at the start and a G at the end). Figure generated using IGV.

**Supplementary Figure 3**. Relationship between *blaTEM* gene copy number and antimicrobial susceptibility. *blaTEM* gene copy number was calculated by dividing its read depth by the average depth of chromosomal genes (*gyrA, gyrB, mefA, parC, acnB, rplD, rplV, mtrR, mtrD, rpsE, rpsJ*) as reported by neissflow, which can also be interpreted as an estimate of plasmid copy number per bacterial chromosome. *blaTEM* copy number was positively associated with penicillin MIC (log2 scale), consistent with a genedosage effect. Associations were modeled using exponential regression with 95% bootstrap confidence intervals; Pearson r (correlation) is reported.

**Supplementary Figure 4:** Graph containing 2 connected components. Nodes are blue circles labeled A-L, edges are black lines, and connected components are denoted via encircling in blue dashed lines (70).

## Supplementary tables

**Supplementary Table S1.** Complete list of AMR associated loci interrogated by neissflow. For each genomic position or amino-acid residue, the table provides the reference (FA19) allele, the default expected state, the corresponding mutation or variant detected, and the associated antimicrobial class. The table includes all nucleotide and amino-acid markers across *23S rRNA, 16S rRNA, penA, mtrR, mtrD, macA, norM, ponA, pilQ, gyrA, parC, gyrB, rpsJ, rpsE, rplD, rplV, ftsX, acnB*, and plasmid-mediated determinants (*blaTEM, tetM*, and macrolide methylases). These represent the full set of AMR determinants evaluated by the neissflow AMR variant analysis script.

**Supplementary Table S2**. Complete List of Validated Analytes with Genetic Basis, Associated Antimicrobials, Standard Methods, and Definitions of True Positive, True Negative, False Positive, and False Negative Classifications for Performance Assessment.

**Supplementary Table S3**: Full neissflow output final report for Dataset 1 (n=158)

**Supplementary Table S4**: Full neissflow output report for Dataset 2 (n=283) for the reproducibility validation (EQA isolates)

**Supplementary Table S5:** Isolate Level Comparison of neissflow’s species detection results to that of the three standard species detection methods used for validation in this study (n=158; Dataset 1)

**Supplementary Table S6**. Isolate Level Comparison of neissflow Detected AMR Markers and Molecular Types with PubMLST and Phenotypic (MIC) Results across 105 Ng isolates (Dataset 1)

**Supplementary Table S7.** Concordance Between neissflow-detected AMR Markers and Antimicrobial Susceptibility Phenotypes.

**Supplementary Table S8.** Reproducibility of neissflow Analyte Calls Across 17 Reference Strains.

**Supplementary Table S9:** Full neissflow output report for isolates with *23S rRNA A2059G* (n = 3) and *C2611T* (n = 25) mutations identified among U.S. national surveillance Ng isolates collected during 2023–2025 (n = 4,084).

**Supplementary Table S10.** Non-*Ng* and low-quality *Ng* samples with <85% FA19 genome coverage at ≥10× depth. Mapping coverage and Mash species-assignment results for samples that failed the *Ng* genome coverage threshold. The table includes non-*Ng* samples and low-quality *Ng* samples with <85% of the FA19 reference genome covered at ≥10× depth. These data support the use of the ≥85% FA19 genome coverage threshold as an internal validation criterion for distinguishing high-quality *Ng* genomes from low-quality or non-*Ng* samples.

