## Supplementary Methods for "neissflow: Streamlining Genomic Epidemiology of *Neisseria gonorrhoeae* with Nextflow"

### Species ID Subworkflow - Mash Hit Determination:

For each sample, the top species hit is selected based on the highest combined identity and hash fraction, excluding plasmid sequences. Additional hits meeting identity ≥ 0.95 and hash fraction ≥ 0.95 but differing from the primary *Ng* species call are reported as potential contaminants, and plasmid sequences meeting these thresholds are also recorded separately.

### AMR Variant Call Evaluation:

To evaluate AMR determinants, the script cross-references variants in the input Snippy VCF file with a curated list of AMR relevant genomic positions and their corresponding reference nucleotides or amino acids from FA19. For each position, the script searches the VCF files for the presence of variants altering nucleotides or amino acids. When no variant is present, the depth of coverage at the position of interest is checked. If the coverage is ≥10x, the script will report that position as containing the reference nucleotide (wild type); if coverage is <10x, the value is reported as "NF" (not found). This is consistent with the default minimum depth of coverage for Snippy to report a variant. The depth of coverage is also used to determine the presence of plasmid AMR genes. If a plasmid AMR gene is found to have a depth of ≥10x, its presence will be reported as "True" and otherwise will be reported as "False". In addition, the VCF files are also used to detect truncations in the *mtrR* and *pilQ* genes, denoted by Snippy with "stop gained". The script also screens for insertions in *rplV* (reported by Snippy as "ins") and for the characteristic duplicate aspartic acid at amino acid position 345 in *penA*, which Snippy reports as "dup". For multicopy 23S rRNA loci, neissflow reports both the nucleotide call and the read-supported allele frequency at resistance-associated positions, including 23S rRNA A2058G, A2059G and C2611T. The reported frequency represents the proportion of mapped reads supporting the variant allele at that position. For example, frequencies near 1.0 suggest that most or all mapped 23S rRNA copies carry the mutation, whereas intermediate frequencies such as ~0.25, ~0.50, or ~0.75 may be consistent with approximately one, two, or three mutated copies, respectively, depending on read mapping and coverage. Therefore, these values should be interpreted as read-supported allele fractions rather than definitive copy-resolved 23S rRNA mutation counts.

**Phylogeny Subworkflow – Implementation & QC:**

For each sample, Snippy produces a VCF and an aligned FASTA against the chosen reference. snippy-core then aggregates the per-sample outputs into a multi-FASTA whole-genome core alignment (core.full.aln). The alignment is sanitized using snippy-clean_full_aln. By option, the reference sequence can be removed before downstream analyses to prevent inclusion as a tip in the tree.

To enable ascertainment bias correction (ASC), monomorphic A/C/G/T counts are computed from the core alignment and used to generate a partition guide that binds the SNP alignment to an ASC DNA model. Maximum-likelihood trees are inferred with RAxML-NG using rapid bootstrapping with automatic MRE (majority-rule extended) stopping and concurrent best-tree search, followed by midpoint rooting with GoTree. The rooted tree is rendered to a PNG with cluster-based tip colors, and an HTML report embeds the figure and summarizes cluster membership.

A phylogeny QC check is performed on the core alignment and tree. Phylogeny QC reports 6 checks 1) number of sequences present in the final WGS alignment: the number of sequences in the core alignment versus the expected sample list; 2) length uniformity: all sequences in the core alignment have identical length; 3) sequence character validity: sequence lines contain only A/C/G/T/N/"-"; 4) tip coverage: all expected samples are present as leaf labels in the Newick file; 5) branch-length outliers: leaves with branch length greater than the mean plus two standard deviations across leaf branch lengths are listed; and 6) core genome size (bp). Except for the branch-length outlier screen and the core genome size, all checks receive a pass/fail designation. The QC table (phylogeny_qc_report.tsv) records, for each check, the accepted value, observed value, and pass/fail status.

**Phylogeny Subworkflow - Outbreak Detection:**

A custom outbreak detection script uses the snp-dists distance matrix to predict outbreak clusters, representing samples as nodes in a graph, with edges defined between pairs with a SNP distance of less than or equal to the input threshold (default: 20 SNPs). The edge creation algorithm is explained by the following piecewise function:

$$f(u,v) = \begin{cases} 1, & \text{SNPdist}(u,v) \leq 20 \\ 0, & \text{otherwise} \end{cases}$$

Eq 1: Piecewise function for edge creation between nodes in outbreak graph (G = (V, E)) where V is the set of all nodes, E is the set of all edges, and u, v ∈ V.

To determine outbreak clusters, the disjoint set union (DSU) algorithm was implemented to identify the connected components in the graph. Connected components are subgraphs within a larger graph in which there exist paths between all the nodes. Supplementary figure 4 gives an example of a graph containing connected components.

The pseudocode for the DSU algorithms Find (Algorithm 1) and Union (Algorithm 2) functions can be found below:

```
Function Find(parent, x) :
  Input: array parent, element index x
  Output: representative (root) of the set containing x

  If parent[x] = x :
    Return x
  Else:
    parent[x] ← Find(parent, parent[x])    // Path Compression
    Return parent[x]
```

**Algorithm 1:** Find function for DSU - Algorithm that finds the representative of the disjoint set that node x is a member of (i.e. the parent of node x). This is a recursive function with a base case of returning the index of the node that is its own parent (the representative of the disjoint set).

```
Function Union(parent, u, v) :
  Input: array parent, elements u, v

  root_u ← Find(parent, u)
  root_v ← Find(parent, v)

  If root_u ≠ root_v :
    parent[root_u] ← root_v    // Link root_u to root_v
```

**Algorithm 2:** Union function for DSU - Algorithm that merges disjoint sets where edges exist between nodes. It uses the Find function to identify disjoint sets by their representative (disjoint sets will have different representatives). The resulting connected components define clusters, which are assigned to deterministic hex colors for tree annotation.

**Analysis 2 – Intra-MLST Phylogenetic Resolution:**

Each analysis was performed using a reference genome from within that MLST group to maximize phylogenetic resolution and detect recent recombination events:

1. MLST 1901 (n=30 isolates): SPL-4 and NG-Y strains; whole-genome alignment generated using NG-Y reference genome (NCBI accession number: CP145017).
2. MLST 7363 (n=67 isolates): NG-X, NG-Z, NG-K, and NG-W strains; whole-genome alignment generated using NG-K reference genome (NCBI accession number: CP145048).
3. MLST 7367 (n=50 isolates): NG-M, NG-U, and CDC 10328 strains; whole-genome alignment generated using NG-U reference genome (NCBI accession number: CP145024)
