## Supplementary Figures for "neissflow: Streamlining Genomic Epidemiology of *Neisseria gonorrhoeae* with Nextflow"

**Supplementary Figure 1:** Phylogenetic validation of neissflow using replicate sequences. (A) Intra-MLST phylogenetic clustering for MLST 1901 (n=30 isolates) from two reference strains (NG-Y, SPL-4), aligned to the NG-Y reference genome. Both strains formed distinct monophyletic clades with 100% accuracy; mean within-strain SNP distance was 0.26, with all pairwise comparisons ≤2 SNPs (see Table 4b). (B) Intra-MLST phylogenetic clustering for MLST 7363 (n=67 isolates) from four reference strains (NG-K, NG-W, NG-X, NG-Z), aligned to the NG-K reference genome. All four strains formed distinct monophyletic clades with 100% accuracy; mean within-strain SNP distance was 0.43, with all pairwise comparisons ≤5 SNPs (see Table 4b). (C) Intra-MLST phylogenetic clustering for MLST 7367 (n=50 isolates) from three reference strains (CDC 10328, NG-M, NG-U), aligned to the NG-U reference genome. All three strains formed distinct monophyletic clades with 100% accuracy; mean within-strain SNP distance was 0.24, with all pairwise comparisons ≤2 SNPs (see Table 4b). Tips are colored by reference strain identity (see legends). All trees were constructed using RAxML-NG with GTR+Γ model and ascertainment bias correction, following recombination removal by Gubbins. Trees are midpoint rooted. The use of MLST-specific reference genomes maximizes phylogenetic resolution and enhances detection of recent recombination events compared to the distantly related FA19 reference used in Figure 4. Figure generated using iTOL (v7.5.1) (62) and Inkscape.

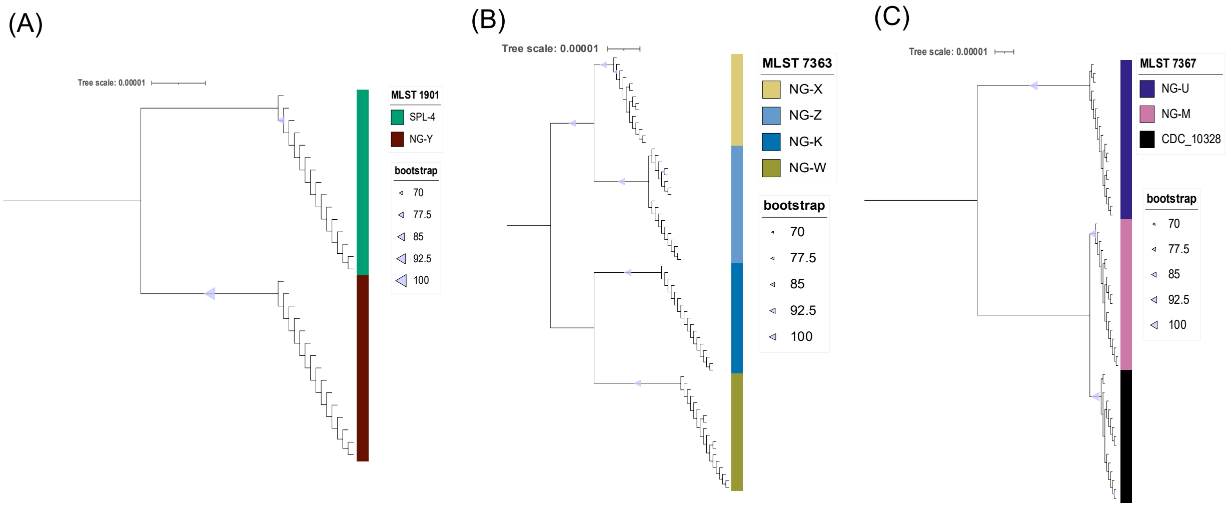


**Supplementary Figure 2:** GCWGS-22115 alignment to FA19 showing the *mtrR* promoter (1110844-1110848) visualized in IGV. This was reported as NFCAAA by neissflow and ACAAG by PubMLST. This deletion would cause the true *mtrR* allele to be delACAA. neissflow parses Snippy output to make this determination. The deletion was reported by Snippy as a complex variant at the second position, resulting in a change from AA to C. In other cases where a deletion has been identified at this position, Snippy has reported a deletion at the position before the start of the *mtrR* promoter sequence (TA to T), and thus an explicitly indicated deletion is what neissflow looks for to report a "del". Even if neissflow had correctly parsed this edge case, the reported variant would have been ACdelAA, since Snippy reports complex variants at the position that they start. This also appears to be inaccurate based on what can be seen at these positions with IGV. PubMLST reports the allele as ACAAG as that is what resolves into the assembly. The G that they are reporting in the last position of the allele is the G from one position over and is not a true SNP (wild type is AAAAA flanked by a T at the start and a G at the end). Figure generated using IGV.

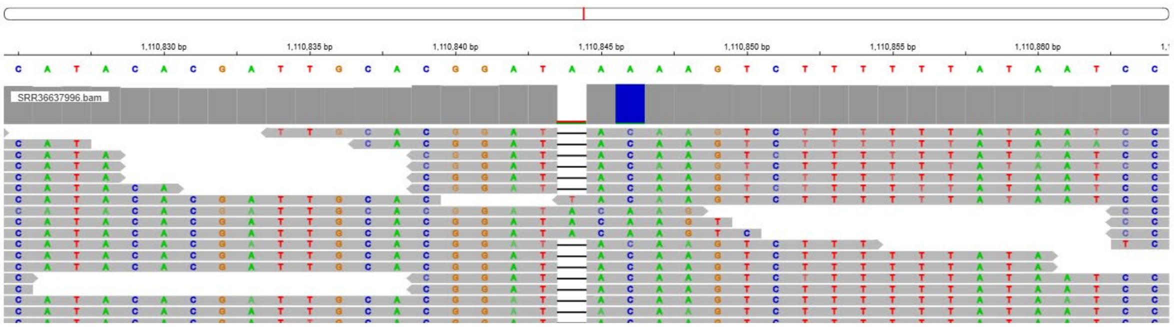


**Supplementary Figure 3**. Relationship between *blaTEM* gene copy number and antimicrobial susceptibility. *blaTEM* gene copy number was calculated by dividing its read depth by the average depth of chromosomal genes (*gyrA, gyrB, mefA, parC, acnB, rplD, rplV, mtrR, mtrD, rpsE, rpsJ*) as reported by neissflow, which can also be interpreted as an estimate of plasmid copy number per bacterial chromosome. *blaTEM* copy number was positively associated with penicillin MIC (log2 scale), consistent with a gene-dosage effect. Associations were modeled using exponential regression with 95% bootstrap confidence intervals; Pearson r (correlation) is reported.

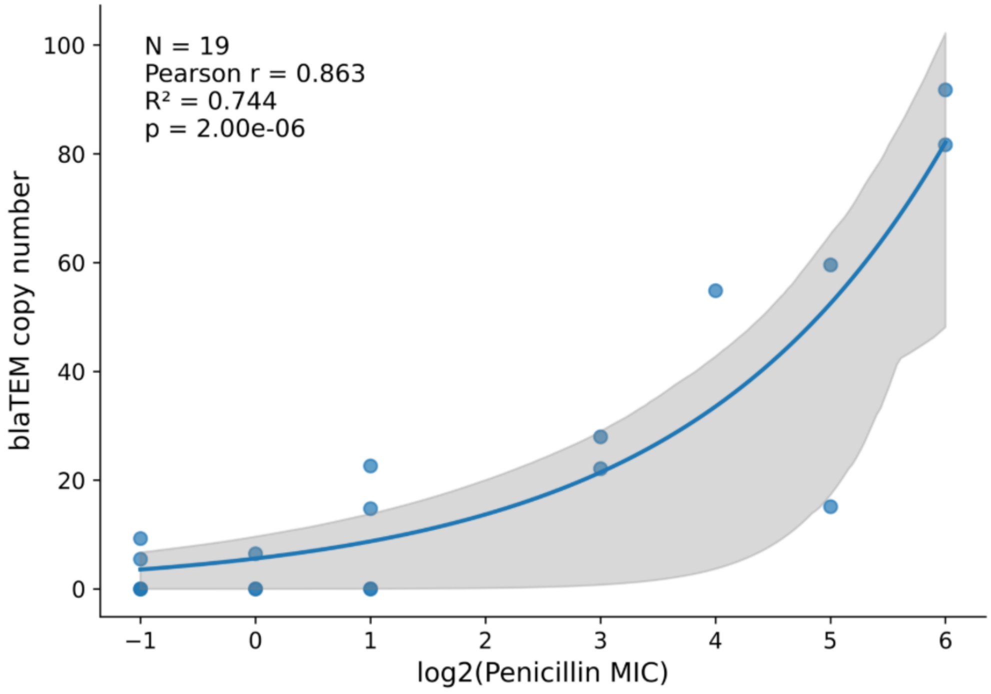


**Supplementary Figure 4:** Graph containing 2 connected components. Nodes are blue circles labeled A-L, edges are black lines, and connected components are denoted via encircling in blue dashed lines (70).

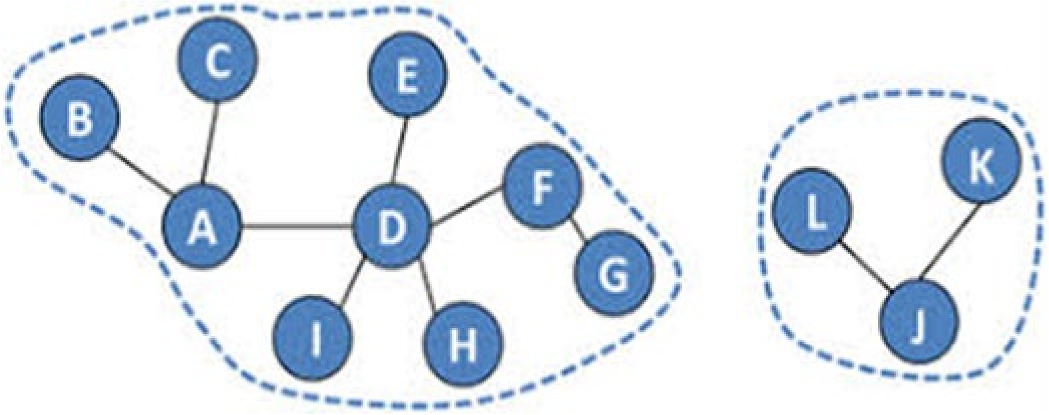
